# Vision Normalizing Flows for the probability-informed detection of banana diseases from in-field images

**DOI:** 10.64898/2026.07.19.739426

**Authors:** Alisa Prusokiene, Augustinas Prusokas, Renata Retkute

## Abstract

Banana diseases impose severe production losses in tropical smallholder farming systems, yet accurate in-field visual diagnosis remains difficult: symptom expression varies across cultivars and growth stages, and several diseases produce morphologically overlapping foliar signs. We developed a probabilistic image-recognition framework for detecting five economically important banana diseases — Xanthomonas Wilt, Banana Bunchy Top Disease, Fusarium Wilt (Panama disease), Yellow Sigatoka, and Black Sigatoka — from in-field photographs, without any disease-specific fine-tuning of the vision backbone. The approach extracts frozen 1,152-dimensional embeddings from the DINOv3 vision foundation model and couples them with a conditional normalizing flow, trained on four publicly available datasets spanning diseased banana plants, healthy tissue, non-banana vegetation, and general natural imagery. On an independent test set the model achieved F1 scores exceeding 0.98, average precision values of 0.968–0.999, and AUROC values of 0.997–1.000 across all five diseases evaluated as binary detection problems. Multi-class accuracy was near-perfect, with limited confusion between Yellow Sigatoka and Black Sigatoka — a biologically plausible ambiguity attributable to overlapping early-infection foliar symptoms. Because the normalizing flow estimates explicit conditional probability densities rather than decision boundaries, two complementary log-likelihood ratios can be derived: a disease ratio comparing each disease class against healthy banana, and a plant ratio comparing banana against non-banana imagery. Together these define an interpretable two-dimensional diagnostic space that simultaneously quantifies evidence for disease presence and image relevance, cleanly separating diseased plants, healthy plants, and out-of-distribution images while flagging uncertain predictions for confirmatory testing. Inference on frozen embeddings is lightweight and compatible with smartphone deployment, providing a scalable, uncertainty-aware diagnostic tool for smallholder farming systems and disease surveillance programmes.

## 1 INTRODUCTION

Bananas and plantains (*Musa* spp.) are among the world’s most important food crops, providing a dietary staple for more than 400 million people and a primary source of income for millions of smallholder farmers across tropical and subtropical regions (Ortiz and Swennen, 2014). Global banana production exceeds 120 million tonnes annually, yet the crop remains genetically narrow and highly vulnerable to emerging and established disease threats (Reay, 2019). Beyond their contribution to food security, bananas support rural livelihoods through local consumption, domestic markets, and international trade (Adenle and Azadi, 2026). Sustaining production is increasingly challenged by plant pathogens whose spread is facilitated by the movement of planting material, climate variability, expanding trade networks, and intensification of agricultural systems (Drenth and Kema, 2021; Bebber, 2019).

Several diseases severely constrain banana production worldwide, including Banana Xanthomonas Wilt (BXW), Fusarium Wilt (Panama disease), Banana Bunchy Top Disease (BBTD), Yellow Sigatoka, and Black Sigatoka (Nakato et al., 2018; Ploetz, 2015; Dita et al., 2018; Churchill, 2011). Together, these diseases cause substantial yield losses, threaten food security, and undermine the livelihoods of smallholder producers. While their causal agents and epidemiology differ, reliable field diagnosis remains difficult because symptom expression varies across cultivars, environmental conditions, disease stages, and management practices (Tripathi et al., 2009; Kumar et al., 2015; George et al., 2022). Early and accurate disease identification is therefore essential for implementing effective control measures, limiting pathogen spread, and reducing economic losses (Premabati and De Mandal, 2020).

The importance of accurate diagnosis extends beyond immediate disease management. Recent epidemiological modelling of BBTV-endemic smallholder landscapes demonstrated that farmer ability to correctly identify disease symptoms is one of the strongest predictors of management adoption and economic outcomes (Retkute et al., 2026). These findings suggest that strengthening disease recognition has the potential to improve not only surveillance effectiveness but also broader livelihood and food-security outcomes.

Traditional disease diagnosis relies primarily on visual scouting by farmers, extension officers, and plant-health specialists. When symptoms are ambiguous or confirmation is required, laboratory testing is typically used. Diagnostic methods include serological assays such as enzyme-linked immunosorbent assays (ELISA) and molecular techniques including polymerase chain reaction (PCR), recombinase polymerase amplification (RPA), and loop-mediated isothermal amplification (LAMP), which detect pathogen proteins or nucleic acids directly (Nakato et al., 2013; Chen and Hu, 2013; Aduo et al., 2025; McFarlane et al., 2026; Ouedraogo et al., 2026). Although these methods provide high sensitivity and specificity, they often require specialised equipment, laboratory infrastructure, and trained personnel, making routine deployment difficult in many smallholder production systems. Visual diagnosis, while inexpensive and accessible, is susceptible to observer bias and frequently fails to distinguish diseases during early infection or when symptoms are atypical. Delayed or incorrect diagnosis consequently hampers timely intervention and facilitates disease spread, highlighting the need for scalable and field-deployable diagnostic tools.

Recent advances in computer vision have created new opportunities for automated disease recognition from digital images (Priyadarshini and Vinothini, 2025). Early approaches to banana disease detection relied on handcrafted features and conventional machine-learning methods (Liao et al., 2019; Ani Brown Mary et al., 2020; Chaudhari and Patil, 2020; Gaur et al., 2023; Cárdenas-Rodríguez et al., 2023). More recently, convolutional neural networks (CNNs) have achieved high classification accuracy across a range of banana diseases, including BXW, Fusarium Wilt, BBTD, and Sigatoka infections (Selvaraj et al., 2019; Sangeetha et al., 2023; Thomas and David, 2023; Sharma et al., 2023; Ashoka S et al., 2024; Shukla et al., 2024; Mora et al., 2025; Banerjee et al., 2025; Genove et al., 2026; Ouedraogo et al., 2026). Despite high reported accuracies, most existing models remain fundamentally discriminative and operate under a closed-set assumption in which every image is assigned to one of the classes observed during training.

This assumption is rarely satisfied in real agricultural environments. Plants may exhibit previously unseen diseases, mixed infections, abiotic stresses, nutrient deficiencies, atypical symptom expression, or image artefacts that are absent from the training data. Under such conditions, conventional classifiers frequently produce highly confident predictions despite limited evidence supporting a particular diagnosis. Consequently, achieving high classification accuracy alone is insufficient for practical deployment; diagnostic systems must also quantify uncertainty and recognise when an image falls outside the distribution represented by the training data.

To address these limitations, there is growing interest in probabilistic, uncertainty-aware approaches capable of identifying samples that deviate from the training distribution, a capability known as out-of-distribution (OOD) detection. Normalizing flows offer a principled solution: by learning invertible mappings between complex data representations and tractable latent distributions, they provide explicit likelihood estimates for any input, enabling both disease classification and anomaly detection within a unified framework (Rezende and Mohamed, 2015; Dinh et al., 2017; Kobyzev et al., 2021). Unlike approximation-based uncertainty methods such as Monte Carlo dropout or deep ensembles, normalizing flows yield exact, interpretable probability estimates and remain computationally efficient at inference time. Despite their theoretical appeal, normalizing flow models have not been systematically evaluated for banana disease detection, and their combination with modern vision foundation models remains unexplored. This study addresses this gap by developing and evaluating a normalizing flow model built on top of a pretrained vision foundation model DINOv3 (Siméoni et al., 2025a) for multi-class banana disease classification and probabilistic diagnostics. While DINOv3 has recently been adopted for leaf lesion segmentation in other crops, notably through integration with UNet architectures for precise delineation of disease regions (Chen et al., 2026; Jiang et al., 2026), its potential for probabilistic, uncertainty-aware disease classification remains largely unexplored. We assess binary and multi-class detection performance across five economically important diseases - BXW, Bunchy Top, Yellow Sigatoka, Black Sigatoka, and Fusarium Wilt - using a large, multi-source field image dataset. We further evaluate the calibration of likelihood-based confidence scores and the system’s capacity to identify out-of-distribution images — capabilities absent from existing closed-set classifiers — and demonstrate their potential for scalable, uncertainty-aware deployment in smallholder farming systems.

## 2 MATERIALS AND METHODS

### 2.1 Model architecture

Our approach combines a pretrained vision foundation model with a conditional normalizing flow to produce probabilistic predictions from in-field banana images (Figure 1). The foundation model extracts high-level semantic image representations that are largely invariant to viewpoint, illumination and background variation, while the conditional normalizing flow models the probability density of these representations conditioned on disease class. This separation of visual feature extraction and probabilistic inference allows the system to exploit the representational power of large-scale self-supervised learning while retaining the calibrated uncertainty estimates afforded by likelihood-based generative models.

**Figure 1.**
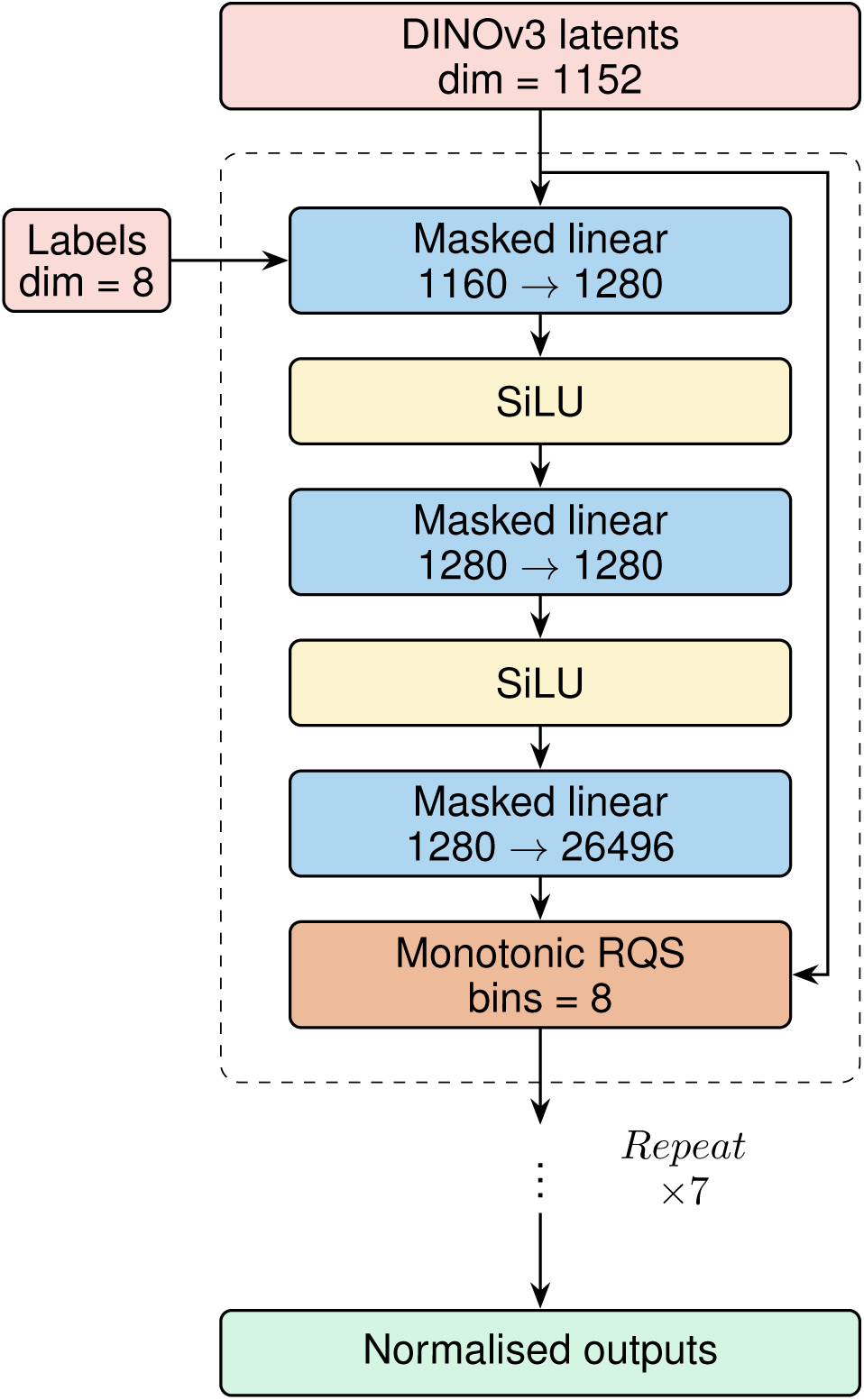
Architecture of the conditional normalizing flow. Frozen 1152-dimensional DINOv3 image embeddings are concatenated with an eight-dimensional conditioning vector and passed through eight autoregressive neural spline transformations. Each transformation comprises masked fully connected networks with SiLU activations that parameterise monotonic rational-quadratic spline bijections, enabling exact likelihood evaluation of the latent representation conditioned on image class.

#### Foundation vision model

Rather than learning visual features directly from the available banana disease images, we adopted a transfer-learning strategy by extracting frozen image embeddings from the pretrained DINOv3 vision foundation model (Siméoni et al., 2025b). DINOv3 is a self-supervised Vision Transformer trained on hundreds of millions of natural images using self-distillation without human annotations, producing representations that transfer effectively to a broad range of downstream computer vision tasks without task-specific fine-tuning.

For an input image, DINOv3 outputs one classification token, four register tokens and a variable number of patch tokens whose number depends on the input image resolution. The classification token summarises global image semantics, the register tokens provide additional persistent latent representations that improve downstream feature learning, while the patch tokens encode local image structure.

To reduce the dimensionality presented to the normalizing flow while retaining information from both global and local image representations, we used the ViT-S variant of DINOv3, which produces embeddings of dimension 384. The four register tokens were averaged to produce a single register representation, and all patch tokens were similarly averaged to obtain a single patch representation. These were concatenated with the classification token to form a fixed-length 1152-dimensional latent vector,

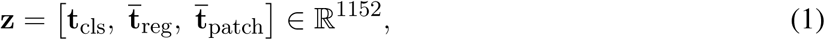

which served as the input to the conditional normalizing flow. Averaging the register and patch tokens substantially reduces computational complexity while preserving the global contextual information required for disease discrimination.

#### Conditional normalizing flow model

The conditional density model was implemented using the PyTorch library Zuko (Rozet et al., 2023), which provides efficient implementations of invertible normalizing flow architectures. Given an image embedding z, the objective of the model is to estimate the conditional probability density

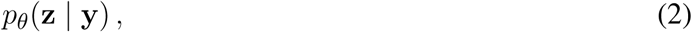

where **y** denotes the disease class conditioning vector and *θ* represents the learnable model parameters.

The flow consists of a sequence of autoregressive neural spline transformations. Each transformation is parameterised by masked fully connected networks employing SiLU activation functions and monotonic rational-quadratic spline (RQS) bijections (Figure 1). Successive transformations progressively map the complex multimodal distribution of DINOv3 embeddings onto a simple multivariate Gaussian latent distribution while preserving exact invertibility. Unlike affine coupling layers, rational-quadratic splines provide substantially greater flexibility for modelling highly non-linear densities without sacrificing analytical tractability.

Class information is incorporated through an eight-dimensional conditioning vector supplied to every autoregressive transformation. At inference time, the same image embedding is evaluated independently under each disease condition to obtain a set of conditional log likelihoods.

Unlike conventional discriminative classifiers, which directly estimate class probabilities through a softmax layer, the normalizing flow explicitly models the distribution of image representations associated with each disease. Classification is therefore performed by selecting the class with the highest conditional likelihood, while the relative differences between likelihoods provide a natural measure of prediction confidence. Furthermore, because the model estimates probability densities rather than only decision boundaries, it readily supports likelihood-ratio tests for binary disease detection and the identification of samples lying outside the training distribution.

### 2.2 Datasets

The training, validation, and test images used in this study were collected from publicly available sources, namely those of (Ouedraogo et al., 2026), (Mduma and Elinisa, 2023), (Garcin et al., 2021), and (Unsplash, 2025).

The first and most varied of the four image datasets, (Ouedraogo et al., 2026), consists of field images that were captured using Android smartphones, quality-filtered, and annotated by expert plant pathologists. The dataset is composed of 17,703 images across 22 disease, stress, and healthy tissue classes. We took particular interest in Banana leaf and whole-plant images affected by Banana Xanthomonas Wilt, Banana Bunchy Top Virus, Yellow/Black Sigatoka, and Fusarium Wilt, as catching these diseases early in the plant’s growth would show the greatest practical benefit. To complement these disease-presenting images, we also included healthy plant images, specifically those of healthy leaves, whole plants, as well as the dried, senescent leaves that naturally form during the plant’s growth cycle.

The second dataset, published by (Mduma and Elinisa, 2023), provides high-quality field images collected from banana plantations in Tanzania. These images substantially increased the representation of Black Sigatoka, Fusarium Wilt and healthy banana plants, improving class balance and increasing the diversity of environmental conditions, cultivars and image acquisition settings encountered during training.

To reduce the likelihood of false-positive disease predictions on unrelated vegetation, we additionally incorporated images from the Pl@ntNet-300K dataset (Garcin et al., 2021). A subset of non-banana plant species was included as a dedicated negative class. This encourages the model to distinguish banana plants from visually similar vegetation before assigning disease labels, thereby improving robustness during deployment in heterogeneous agricultural environments.

Finally, we incorporated a large collection of general natural images from the Unsplash Lite dataset (Unsplash, 2025). These images contain landscapes, people, buildings, animals and miscellaneous objects unrelated to banana cultivation. Inclusion of this class exposes the model to a substantially broader image distribution than agricultural datasets alone and provides an effective out-of-distribution reference during training.

Images of the same class, such as those that exhibit the same disease or of healthy plants, were pooled across the datasets in order to create a single, combined image datasets. The contribution of each source dataset to a given class is shown in Table 1.

**Table 1.** Contributions of banana and non-banana image datasets to each label class.

| Class | Dataset | Size |
| --- | --- | --- |
| Xanthomonas Wilt | Ouedraogo et al. (2026) | 408 |
| Bunchy Top | Ouedraogo et al. (2026) | 980 |
| Yellow Sigatoka | Ouedraogo et al. (2026) | 1, 590 |
| Black Sigatoka | Mduma and Elinisa (2023) | 5, 208 |
| Black Sigatoka | Ouedraogo et al. (2026) | 1, 311 |
| Fusarium Wilt | Mduma and Elinisa (2023) | 3, 944 |
| Fusarium Wilt | Ouedraogo et al. (2026) | 988 |
| Healthy | Mduma and Elinisa (2023) | 4, 888 |
| Healthy | Ouedraogo et al. (2026) | 1, 227 |
| Plantnet | Garcin et al. (2021) | 50, 301 |
| Unsplash Lite | Unsplash (2025) | 24, 996 |

Class sizes were heavily imbalanced, with the largest (Unsplash Lite) containing 24,996 images, and the smallest (Xanthomonas Wilt) being comprised of just 408, which required careful sampling to avoid performance degradation during training.

### 2.3 Image preprocessing

#### Image augmentation

To increase the size and diversity of training images, especially in the case of the smaller classes, we utilised the Python image augmentation library AlbumentationsX (Buslaev et al., 2025). The full augmentation procedure is given in Figure 2. To preserve performance on unaugmented images, we made sure to keep at least one original copy of each image.

**Figure 2.**
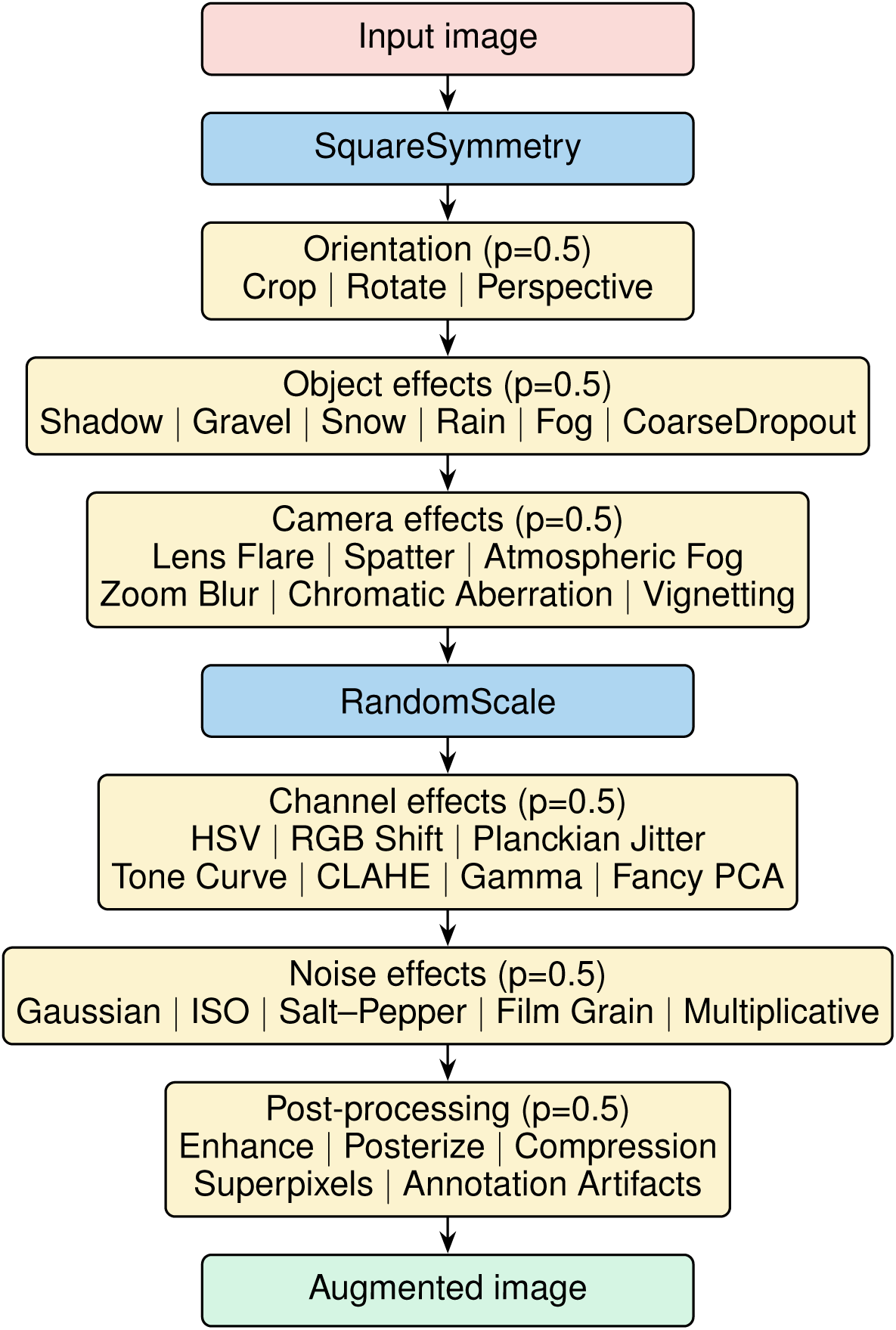
Image augmentation pipeline applied during training. Every image underwent symmetry augmentation and random scaling, followed by stochastic selection of at most one transformation from each augmentation category. These categories comprised geometric transformations, simulated environmental effects, camera artefacts, colour perturbations, image noise and post-processing operations. At least one unmodified copy of every original image was retained within the training set.

Final training, validation, and test dataset sizes are given in Table 2. Additionally, to prevent information leakage between the training, validation and test sets, the dataset was partitioned into these subsets prior to image augmentation. All augmented versions of a given image inherited the subset assignment of the original image from which they were generated, ensuring that no augmented variants of the same source image appeared in multiple data partitions.

**Table 2.** Train-validation-test split for each image class. Training images were augmented using the AlbumentationsX library to increase the size of under-represented classes. Dataset partitioning was performed prior to augmentation, and all augmented images remained within the same partition as their corresponding original image to prevent data leakage between the training, validation and test sets.

| Class | Train |  |  | Validation | Test |
| --- | --- | --- | --- | --- | --- |
|  | Original | Augmented | Total | Original | Original |
| Xanthomonas Wilt | 320 | 31,680 | 32,000 | 43 | 44 |
| Bunchy Top | 776 | 38,024 | 38,800 | 101 | 102 |
| Yellow Sigatoka | 1,272 | 24,168 | 25,440 | 159 | 159 |
| Black Sigatoka | 5,208 | 46,872 | 52,080 | 655 | 656 |
| Fusarium Wilt | 3,944 | 35,496 | 39,440 | 494 | 494 |
| Healthy | 4,888 | 43,992 | 48,880 | 613 | 614 |
| Plantnet | 40,240 | - | 40,240 | 5,030 | 5,031 |
| Unsplash Lite | 19,992 | - | 19,992 | 2,502 | 2,502 |

#### Vision model preparation

Prior to feature extraction, all images were converted to RGB colour space and resized according to the input requirements of DINOv3 while preserving their aspect ratio. Images were normalised using the preprocessing pipeline accompanying the pretrained foundation model. No task-specific fine-tuning of DINOv3 was performed; instead, frozen image embeddings were extracted once and cached for subsequent normalizing flow training. This substantially reduced computational cost by eliminating repeated forward passes through the vision transformer during optimisation.

### 2.4 Training regimen

To improve generalisation and reduce overfitting, optimisation was performed on blended latent representations rather than individual samples. Each training batch consisted of 4096 latent vectors generated using Dirichlet Mixup. Mixture coefficients were sampled from an eight-dimensional Dirichlet distribution with concentration parameter *α* = 0.1, after which randomly selected latent vectors from each class were linearly combined according to the sampled weights.

The corresponding Dirichlet coefficients were simultaneously supplied as conditioning variables to the normalizing flow. The model therefore learned smooth probability densities over interpolated latent representations rather than isolated observations, encouraging locally consistent likelihood estimates throughout the feature space.

Training minimised the average conditional negative log likelihood,

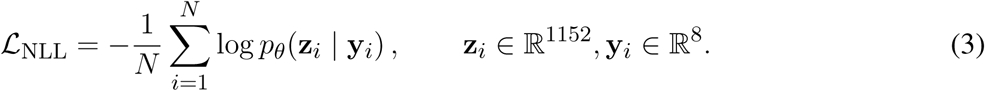

where **z***_i_* denotes the DINOv3 latent representation, **y***_i_* the corresponding conditioning vector, and *N* the batch size. Optimisation continued while monitoring validation negative log likelihood together with downstream classification metrics to identify the point of best generalisation.

Optimisation was performed using the Prodigy optimiser together with the Schedule-Free training framework (Booker, 2024). Prodigy (Mishchenko and Defazio, 2023) is a parameter-free adaptive optimisation method that automatically estimates an appropriate effective learning rate during training, thereby removing the need for extensive manual hyperparameter tuning while maintaining rapid convergence across a wide range of deep learning problems. To further simplify optimisation, we adopted the Schedule-Free (Defazio et al., 2024) approach, which eliminates conventional learning-rate schedules by maintaining two sets of model parameters: one used for optimisation and another averaged set used for evaluation. This approach provides the stability typically associated with learning-rate decay or exponential moving averages without requiring manually designed schedules or warm-up phases. The combination of Prodigy and Schedule-Free proved particularly well suited to optimisation of the conditional normalizing flow, allowing training to converge smoothly while reducing the number of optimisation hyperparameters requiring empirical tuning.

### 2.5 Compute platform

Model development and training were performed using Google Cloud Platform Tensor Processing Units (TPUs). DINOv3 feature extraction and normalizing flow optimisation were implemented in PyTorch, with mixed-precision arithmetic used where appropriate to maximise computational efficiency. Precomputing DINOv3 embeddings substantially reduced training time and enabled optimisation to be performed entirely within latent space.

## 3 RESULTS

We present a machine-learning pipeline for banana disease detection from in-field images, integrating a frozen DINOv3 foundation model with a conditional normalizing flow. We first describe the training and convergence results of the flow, then assess the multi-class classification accuracy of the combined framework, and finally validate its utility as a robust diagnostic system through probabilistic likelihood ratios.

### 3.1 Model training

Frozen DINOv3 embeddings were generated for every image in the training, validation and testing datasets prior to optimisation. By training exclusively within the latent feature space, optimisation became computationally efficient while retaining the rich semantic representations learned by the vision foundation model. Representative training images from the five banana disease classes and the non-disease classes are presented in Figures 3 and 4.

**Figure 3.**
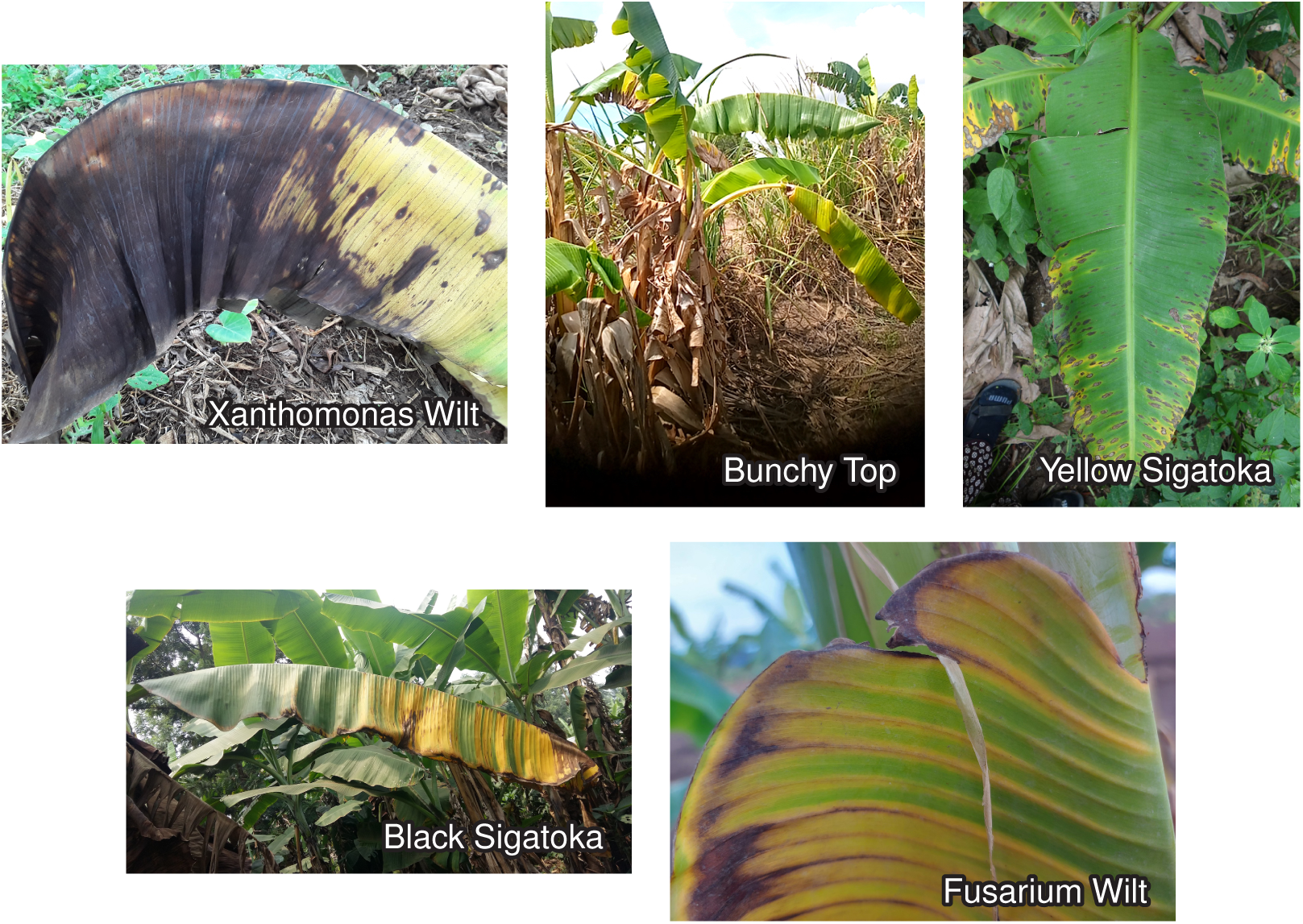
Representative images from the five banana disease classes used during model development. Images illustrate the substantial variation in symptom severity, lighting conditions, plant growth stage, viewing angle and field background encountered during in-field image acquisition.

**Figure 4.**
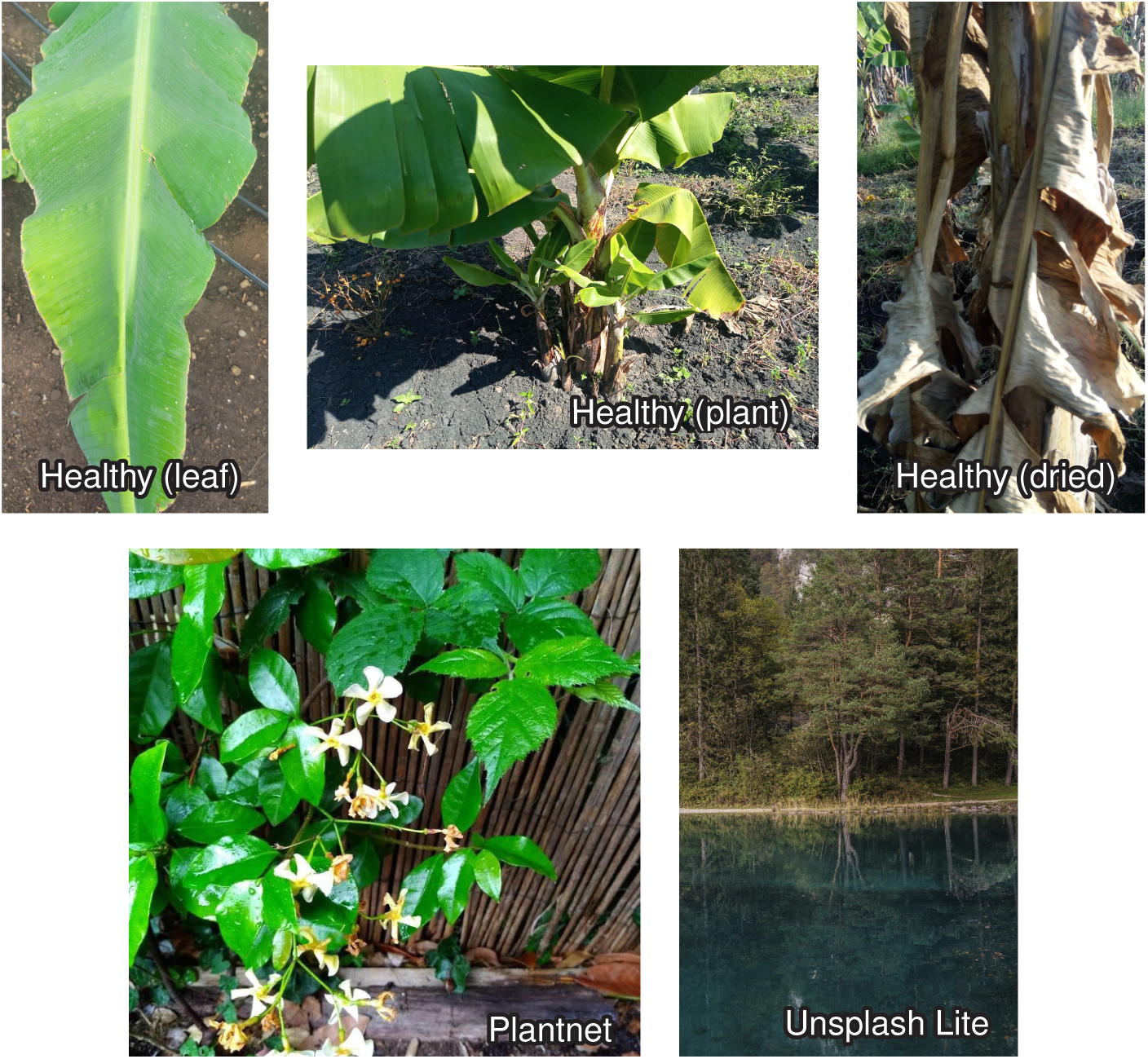
Representative images from the non-disease classes. Healthy banana leaves, healthy whole plants and naturally senescent leaves were included alongside non-banana vegetation (PlantNet) and general natural images (Unsplash Lite) to improve discrimination between diseased banana plants, healthy plants and unrelated scenes.

Throughout training, both density-estimation and downstream classification performance were monitored using the validation set. Figure 5 shows the evolution of average negative log likelihood, macro F1 score, average precision and Area Under the Receiver Operating Characteristic (AUROC) during optimisation. Negative log likelihood decreased rapidly during the early stages of training before gradually converging, while all classification metrics increased correspondingly and stabilised close to their maximum values.

**Figure 5.**
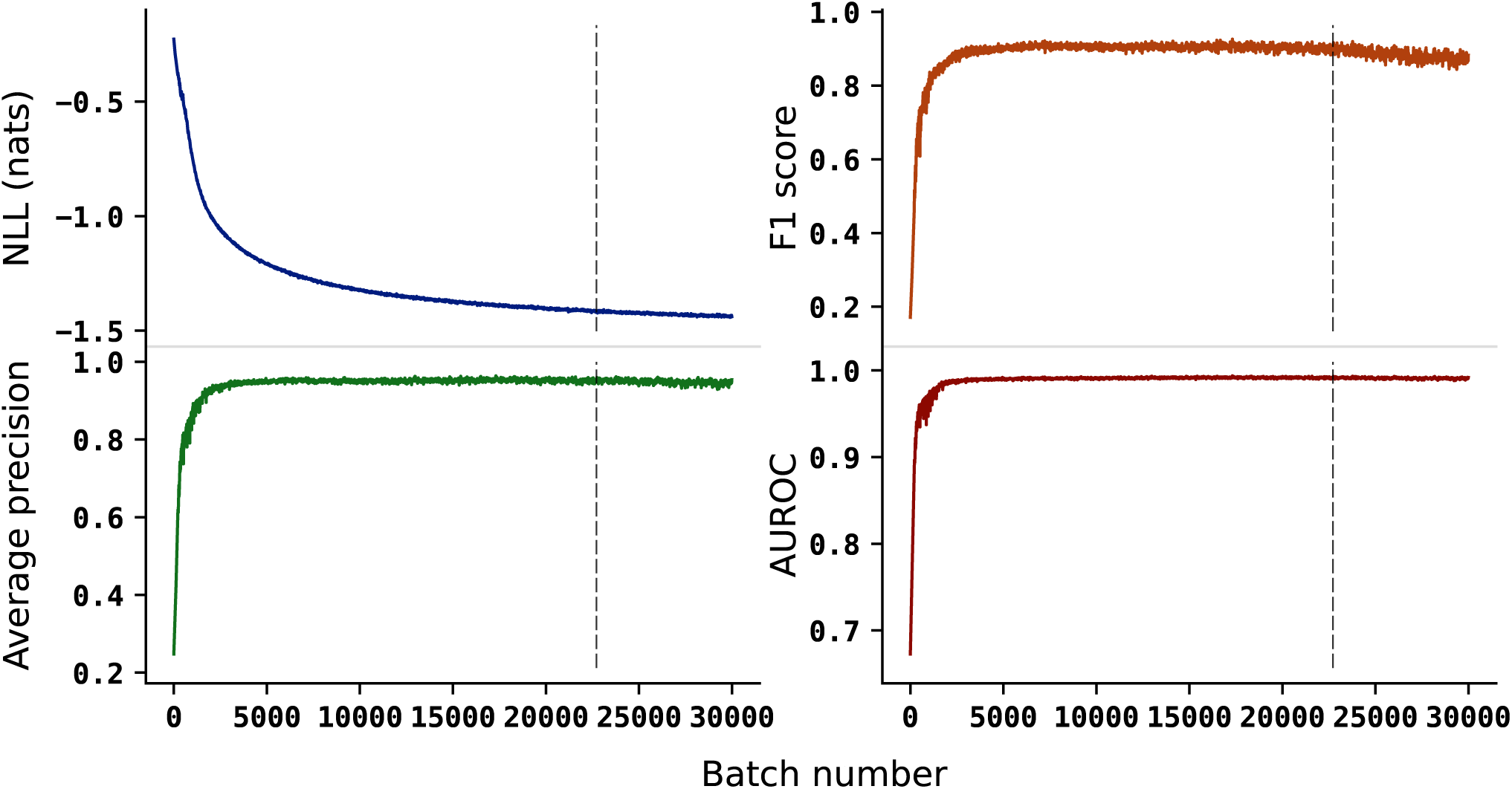
Validation statistics monitored during optimisation. Mean negative log likelihood decreased steadily throughout training, while macro F1 score, average precision and AUROC increased before reaching stable plateaus. The vertical dashed line indicates the checkpoint selected for the final model (Batch `22715`).

The final model checkpoint was selected according to validation performance rather than the final optimisation iteration, providing the best balance between likelihood estimation accuracy and generalisation to unseen images. No evidence of overfitting was observed, with validation performance remaining stable after convergence.

### 3.2 Image classification

Classification performance was evaluated using the independent test dataset comprising all eight image classes: Xanthomonas Wilt, Bunchy Top, Yellow Sigatoka, Black Sigatoka, Fusarium Wilt, Healthy, Plantnet and Unsplash Lite. Predicted labels were assigned according to the maximum conditional log likelihood across the candidate classes.

The resulting confusion matrix (Figure 6) demonstrates high overall classification performance across both disease and non-disease categories. Most disease classes exhibit very few misclassifications, with particularly strong performance observed for Fusarium Wilt, Bunchy Top and Xanthomonas Wilt. Healthy banana plants and the two negative classes (PlantNet and Unsplash Lite) were also recognised with high accuracy, indicating that the model effectively distinguishes banana diseases from unrelated vegetation and general natural imagery.

**Figure 6.**
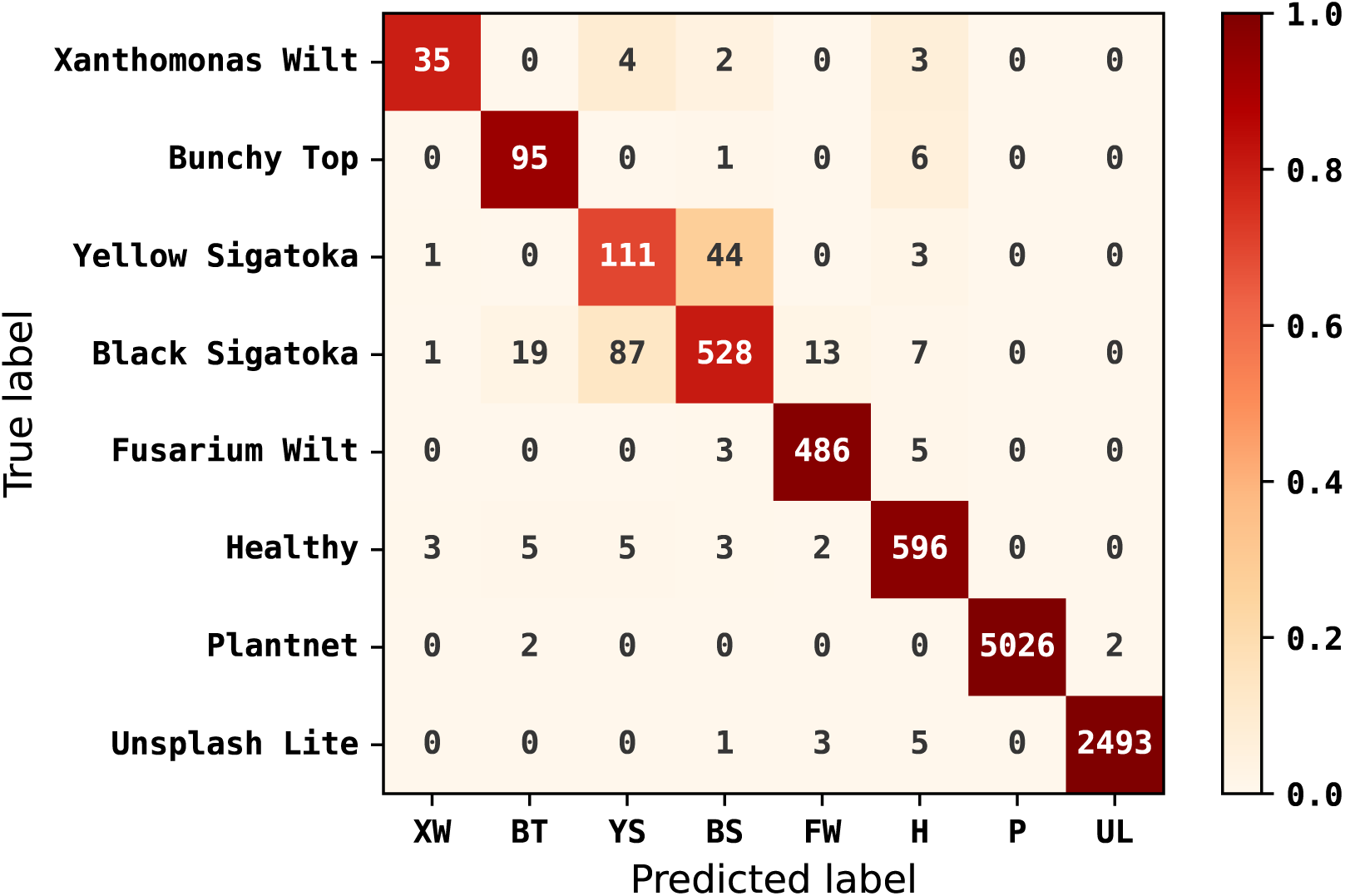
Confusion matrix on the independent test set. Rows correspond to true image classes and columns to predicted classes. Classification was performed by selecting the class with the highest conditional log likelihood.

Confusion between Yellow and Black Sigatoka represented the primary classification error, a biologically plausible ambiguity given the overlapping colour, size, and spatial distribution of foliar lesions during intermediate infection. Nonetheless, the confusion rate remained low relative to the large class sizes, and misclassified images exhibited substantially narrower likelihood margins than correct predictions, confirming that the flow retains informative uncertainty even under ambiguous symptom expression.

Figure 7 illustrates the distribution of classification log likelihood ratios between the highest and second-highest scoring classes. Correct predictions typically exhibited large positive likelihood ratios, reflecting high model confidence, whereas misclassified samples clustered close to zero where competing disease hypotheses possessed similar likelihoods. These results suggest that likelihood ratios provide a useful confidence measure for practical deployment by identifying cases requiring additional inspection.

**Figure 7.**
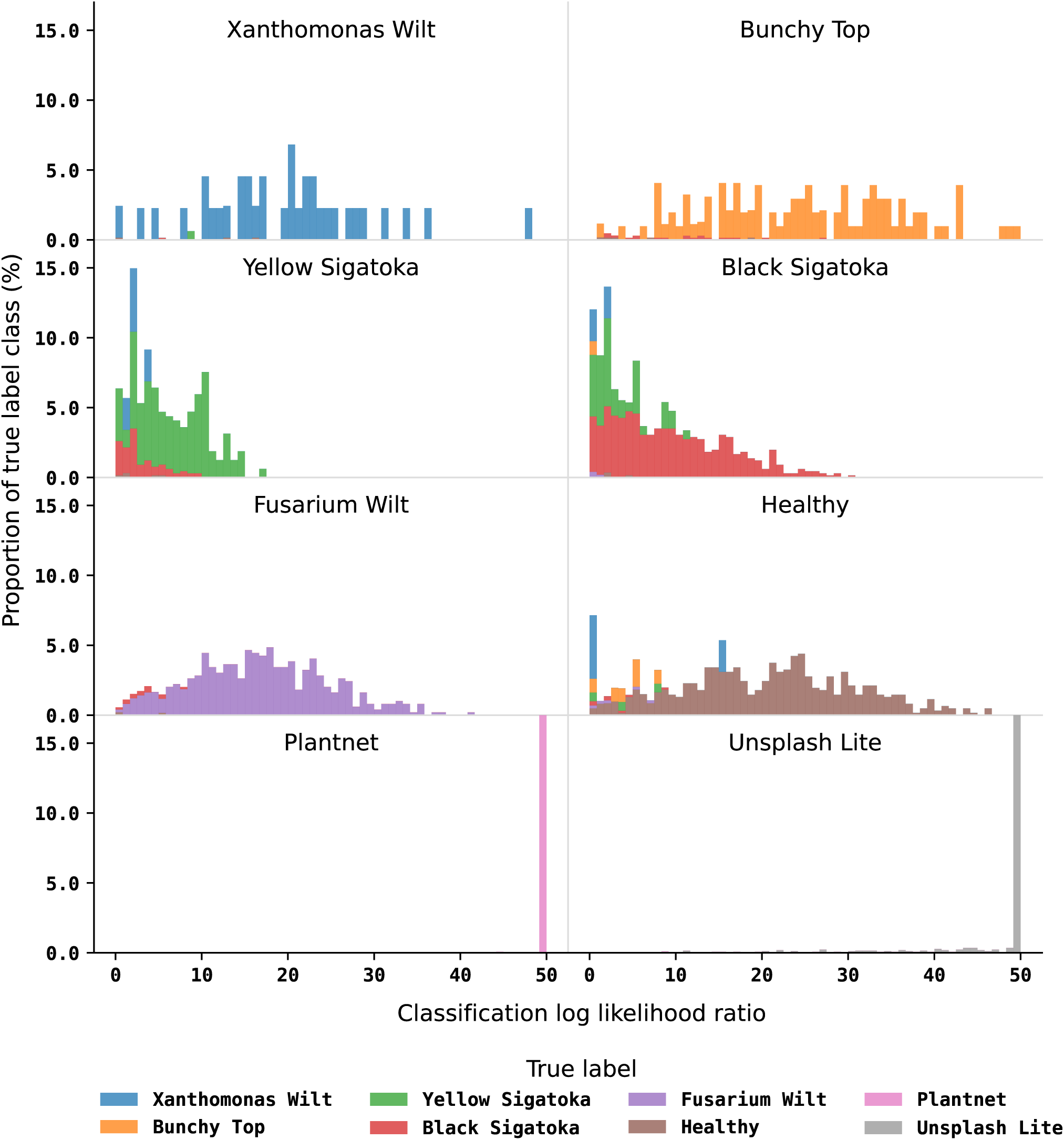
Distribution of classification confidence measured by the log likelihood ratio between the highest- and second-highest-scoring classes. Larger values indicate greater confidence in the predicted label. Histograms are normalised within each true class to facilitate comparison across classes of different sizes, and log likelihood ratio values are clipped to 50.

Examination of misclassified images revealed several recurring failure modes (Figures 8 and 9). The majority of false positives involved healthy banana plants exhibiting natural senescence, environmental stress or complex illumination patterns that partially resembled disease symptoms. A small number of non-banana images containing elongated leaves or strong linear textures also received moderate disease likelihoods.

**Figure 8.**
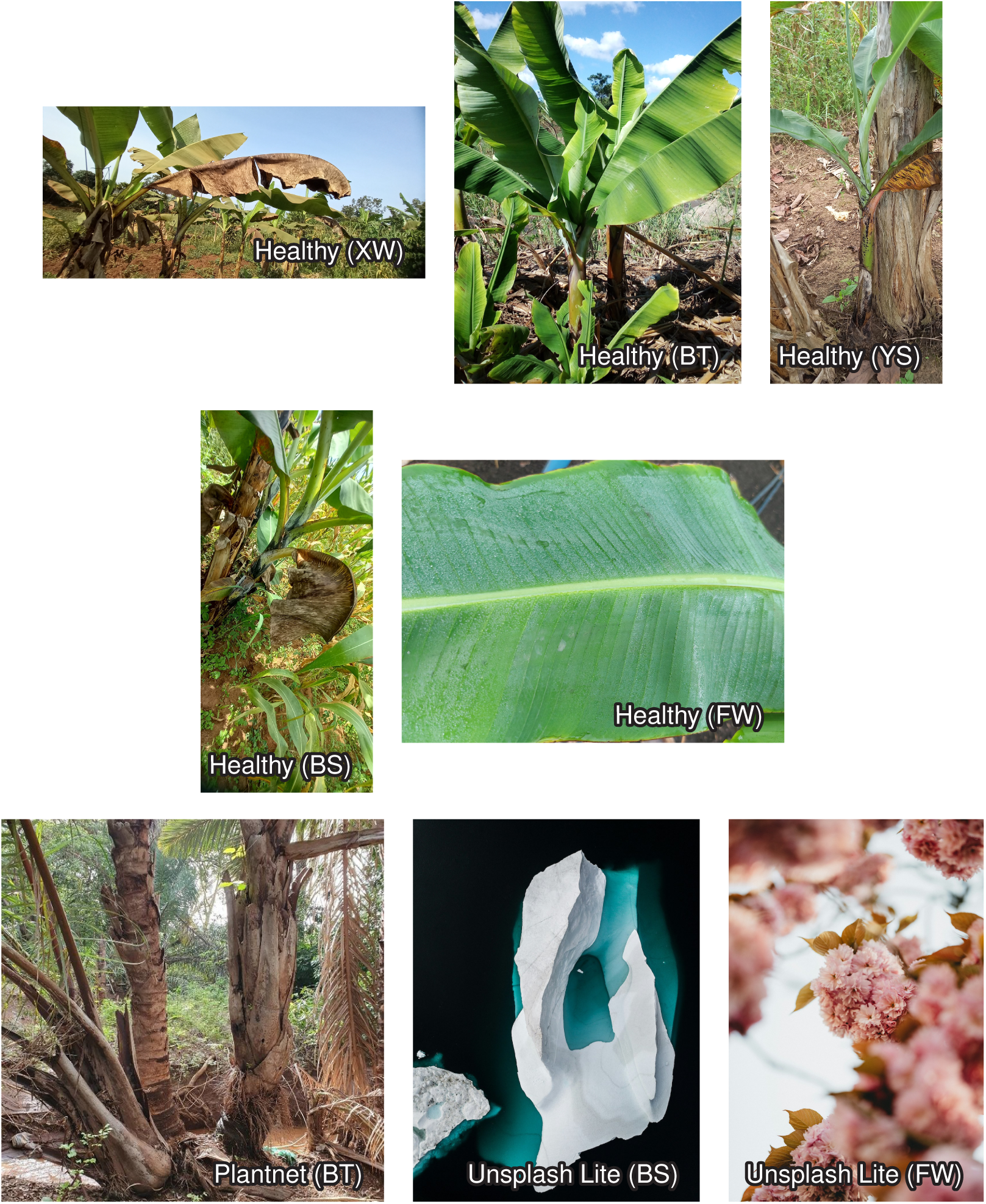
Representative false-positive predictions. Incorrectly predicted labels are given in brackets: Xanthomonas Wilt (XW), Bunchy Top (BT), Yellow Sigatoka (YS), Black Sigatoka (BS), Fusarium Wilt (FW).

**Figure 9.**
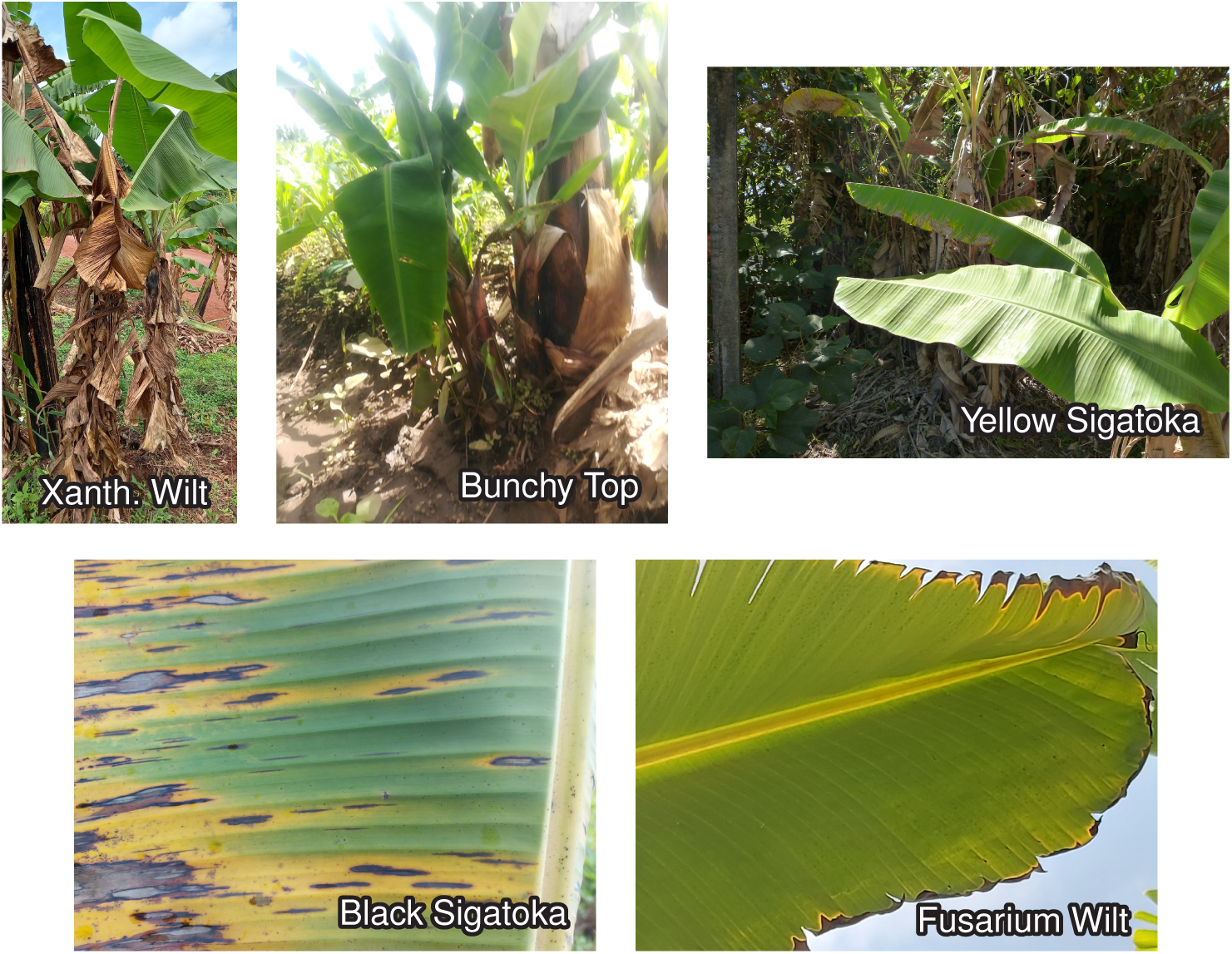
Representative false-negative predictions, illustrating various visual scenarios responsible for missed detections under field conditions.

False negatives generally corresponded to images displaying subtle or atypical symptoms, severe occlusion, poor image quality or mixed symptom presentations. In particular, early-stage Sigatoka infections occasionally lacked sufficiently distinctive visual characteristics for reliable separation from healthy tissue or the alternate Sigatoka class. These observations are consistent with the known biological variability of foliar disease expression under field conditions.

Importantly, even many incorrectly classified images exhibited relatively small likelihood differences between competing classes, suggesting that uncertainty estimates produced by the normalizing flow remain informative even when the highest-scoring class is incorrect.

### 3.3 Disease detection

Beyond conventional multi-class classification, the probabilistic formulation of the normalizing flow enables likelihood ratios to be combined into biologically meaningful diagnostic scores. Two complementary quantities were considered: a banana disease likelihood ratio comparing diseased and healthy banana classes, and a banana plant likelihood ratio comparing banana images against non-banana imagery.

Figure 10 demonstrates clear separation between diseased banana plants, healthy banana plants and unrelated images using these likelihood ratios. Diseased samples consistently produced strongly positive disease likelihood ratios, whereas healthy banana plants clustered around negative values. Similarly, banana images were readily separated from non-banana vegetation and general natural images using the plant likelihood ratio.

**Figure 10.**
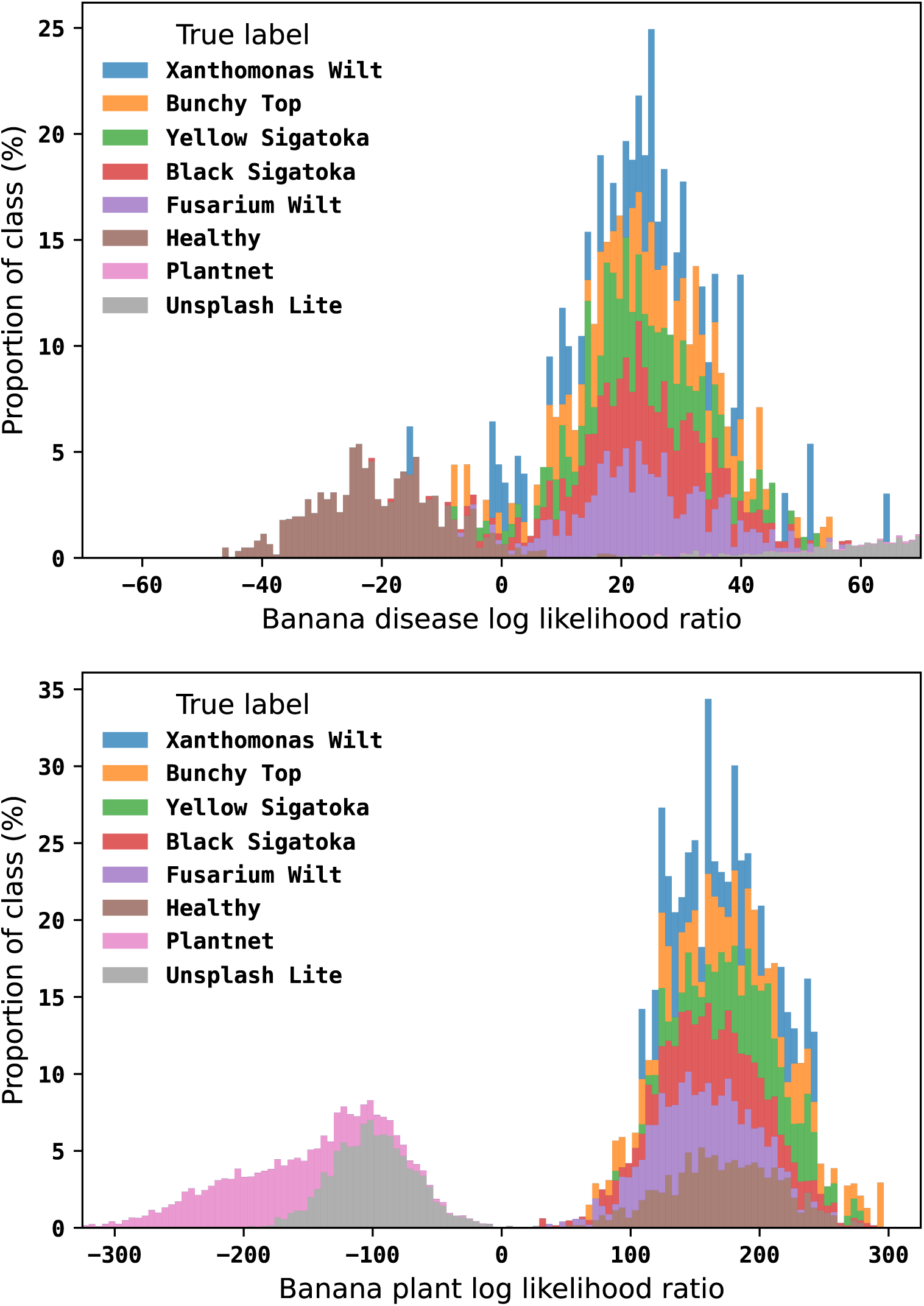
One-dimensional likelihood ratios derived from the conditional normalizing flow. Top: log likelihood ratio comparing disease classes with the healthy banana class. Bottom: log likelihood ratio comparing banana classes with the two non-banana reference classes. Positive values indicate stronger evidence for the numerator hypothesis.

To quantify this capability, each disease was additionally evaluated as an independent binary detection problem by comparing the disease class against healthy banana plants. Performance metrics are summarised in Table 3. Across all five diseases, the model achieved F1 scores exceeding 0.98, with average precision values ranging from 0.968 to 0.999 and AUROC values between 0.997 and 1.000. Fusarium Wilt achieved the highest overall performance (F1 = 0.9927, AUROC = 0.9998), while Xanthomonas Wilt exhibited slightly lower average precision, reflecting its comparatively small training set. Nevertheless, all disease classes demonstrated excellent discriminative performance, indicating that likelihood-based binary detection remains highly reliable even when disease prevalence and class balance differ substantially.

**Table 3.** Binary classification performance of the flow model for each disease class. Test images from a given class, alongside healthy banana plant images, were scored by the model. The F1 metric is calculated from the binarised difference in scores, while Average Precision (AvPr) and Area Under Receiver Operator Characterising (AUROC) are derived from their log likelihood ratios.

| Class | F1 | AvPr | AUROC |
| --- | --- | --- | --- |
| Xanthomonas Wilt | 0.9878 | 0.9678 | 0.9967 |
| Bunchy Top | 0.9846 | 0.9884 | 0.9980 |
| Yellow Sigatoka | 0.9858 | 0.9969 | 0.9991 |
| Black Sigatoka | 0.9818 | 0.9982 | 0.9978 |
| Fusarium Wilt | 0.9927 | 0.9998 | 0.9998 |

Combining the banana disease and banana plant likelihood ratios produced a two-dimensional diagnostic space (Figure 11) that simultaneously measures disease evidence and image relevance. Images lying within the upper-right quadrant correspond to highly confident disease detections, whereas samples located close to either decision boundary exhibit greater uncertainty and may benefit from additional inspection or laboratory confirmation. Unlike conventional softmax classifiers, this representation provides interpretable probabilistic evidence for both disease presence and image validity, making it well suited to practical deployment where previously unseen plant species, environmental artefacts or non-agricultural images may be encountered.

**Figure 11.**
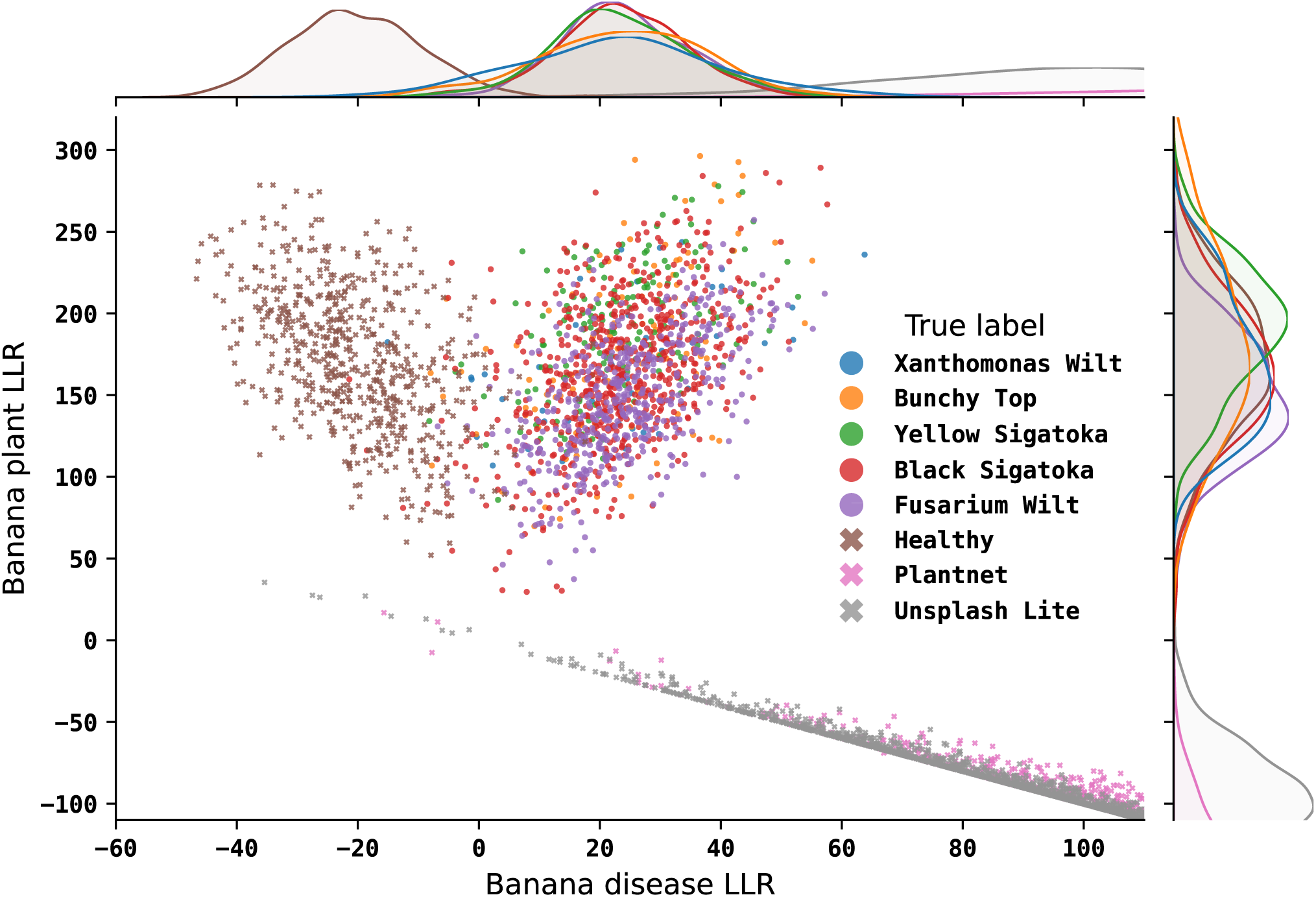
Joint representation of banana disease and banana plant log likelihood ratios. Combining the two probabilistic scores improves discrimination between diseased banana plants, healthy banana plants and unrelated images while providing an interpretable measure of prediction confidence suitable for uncertainty-aware field diagnosis.

## 4 DISCUSSION

This study demonstrates that combining a vision foundation model with a conditional normalizing flow provides an effective framework for probabilistic banana disease diagnosis from field images. Across five economically important banana diseases, the model achieved consistently high classification performance while simultaneously producing interpretable measures of prediction confidence and image relevance. Unlike conventional image classifiers, which typically return only a predicted label, the proposed framework explicitly models class-conditional probability densities and therefore supports uncertainty-aware decision making alongside disease recognition. These results suggest that likelihood-based approaches can complement recent advances in foundation models and address a key limitation of existing image-based diagnostic systems: their inability to recognise when predictions are uncertain or when an image lies outside the training distribution.

A notable finding is that strong diagnostic performance was achieved without disease-specific fine-tuning of the visual backbone. The DINOv3 foundation model was used solely as a frozen feature extractor, yet the resulting system successfully distinguished among BXW, BBTD, Fusarium Wilt, Yellow Sigatoka, Black Sigatoka, healthy banana plants and non-banana imagery. This finding highlights the transferability of modern self-supervised visual representations and suggests that foundation models may substantially reduce the dependence on large disease-specific image datasets. Developing annotated datasets remains one of the primary bottlenecks for machine-learning applications in agriculture, particularly for neglected crops and diseases where data collection and expert labelling are expensive. The performance observed here indicates that much of the information required for disease discrimination is already captured within general-purpose visual representations, allowing the downstream model to focus on learning disease-specific probability distributions rather than low-level image features.

The principal methodological contribution of this study lies in the use of conditional normalizing flows as a diagnostic layer. Most existing crop disease recognition systems are discriminative classifiers that assign every image to one of the predefined classes encountered during training. Such systems often remain highly confident even when presented with unfamiliar symptoms, image artefacts or unrelated objects. By contrast, the normalizing flow explicitly estimates conditional likelihoods for each disease class, providing information about both classification and confidence. Images associated with incorrect predictions generally exhibited much smaller likelihood differences between competing classes than correctly classified images, indicating that uncertainty estimates remained informative even when the top-ranked prediction was incorrect. This behaviour is particularly valuable in practical disease surveillance settings, where identifying uncertain cases may be as important as achieving high average classification accuracy.

An additional advantage of the probabilistic framework is the ability to construct biologically meaningful diagnostic scores through likelihood ratios. Rather than relying on a single categorical prediction, the model produces evidence both for disease presence and for whether an image is likely to represent a banana plant. These complementary measures define a two-dimensional diagnostic space that separates diseased plants, healthy plants and unrelated imagery while naturally identifying ambiguous observations. Such a representation provides a more nuanced interpretation of model outputs than conventional softmax classifiers and may prove particularly useful for field deployment. Images occupying regions of low confidence or conflicting evidence could be flagged for expert review or laboratory confirmation, thereby reducing the risk of inappropriate management decisions arising from overconfident predictions.

The observed classification errors provide further insight into the strengths and limitations of image-based disease diagnosis. The dominant source of confusion involved Yellow Sigatoka and Black Sigatoka, which is biologically plausible given the substantial overlap in lesion appearance during intermediate stages of disease development. This pattern is consistent with findings from Ouedraogo et al. (2026), who reported analogous confusion between these two classes: Black Sigatoka achieved 79.3% accuracy, with 19.2% of samples misclassified as Yellow Sigatoka, while Yellow Sigatoka reached 72.1% accuracy, with 27.1% misclassified as Black Sigatoka. Both diseases are characterised by elongated necrotic lesions that vary considerably with disease progression, environmental conditions and image acquisition angle. The fact that most errors occurred between symptomatically related diseases suggests that misclassifications were driven primarily by genuine visual ambiguity rather than a failure to learn disease-specific features. Similarly, several false positives involved healthy plants exhibiting senescence, environmental stress or unusual illumination patterns that partially resembled disease symptoms. These cases highlight an inherent challenge in field-based diagnosis: symptom expression exists along a continuum rather than as discrete categories, and even expert observers may disagree when confronted with atypical or transitional phenotypes.

The inclusion of non-banana vegetation and general natural imagery proved important for improving robustness. Agricultural image classifiers are often evaluated only on disease classes of interest, despite real-world deployment requiring discrimination between relevant and irrelevant images. The strong performance observed for both PlantNet and Unsplash images suggests that exposure to diverse negative examples allowed the model to learn a broader representation of what does not constitute a banana disease. This capability is particularly important for smartphone-based applications, where users may inadvertently photograph non-target plants, background vegetation or unrelated objects. The ability to identify such images as outside the target domain represents an essential prerequisite for reliable operational deployment.

From an applied perspective, the proposed framework has potential value for disease surveillance and decision support in smallholder farming systems. Accurate field diagnosis is frequently a limiting factor in disease management because laboratory confirmation is often expensive, geographically inaccessible and time-consuming. An uncertainty-aware image-recognition system could support extension officers, surveillance programmes and farmers by providing rapid preliminary assessments while highlighting cases that warrant further investigation. Rather than replacing molecular diagnostics, such systems may help prioritise scarce diagnostic resources and increase surveillance coverage. This may be particularly relevant for diseases such as BBTD and BXW, where early detection and timely intervention are critical for reducing onward spread.

Several limitations should be acknowledged. First, evaluation was conducted using publicly available datasets rather than prospective deployment under operational field conditions. Although the datasets encompassed substantial variation in symptom severity, imaging conditions and environmental backgrounds, performance may differ when applied to new geographic regions, cultivars or farming systems. Second, the framework relies on visible symptom expression and therefore cannot detect latent or asymptomatic infections. Third, mixed infections (Blomme et al., 2026), novel diseases and abiotic stress conditions remain underrepresented within current training data. These factors represent important sources of uncertainty in real agricultural environments and warrant further investigation. Finally, while likelihood-based methods provide valuable confidence information, the relationship between likelihood values and decision thresholds will require careful calibration for specific deployment scenarios.

Future work should focus on prospective validation under field conditions, integration with mobile diagnostic platforms, and expansion towards broader surveillance tasks involving multiple crop species and disease complexes. Combining image likelihoods with complementary information such as geographic location, cultivar identity, environmental conditions or temporal observations may further improve diagnostic reliability. More broadly, the results presented here suggest that coupling foundation-model representations with probabilistic generative modelling offers a promising direction for agricultural artificial intelligence, providing not only highly accurate predictions but also transparent measures of certainty that are essential for trustworthy decision support.

## CONFLICT OF INTEREST STATEMENT

The authors declare that the research was conducted in the absence of any commercial or financial relationships that could be construed as a potential conflict of interest.

## AUTHOR CONTRIBUTIONS

A. Prusokiene: Conceptualization, Data curation, Formal Analysis, Investigation, Methodology, Software, Validation, Visualization, Writing – original draft, Writing – review and editing. A. Prusokas: Conceptualization, Writing – original draft, Writing – review and editing. R. Retkute: Conceptualization, Project administration, Writing – original draft, Writing – review and editing.

## FUNDING

R.R. acknowledge the Mastercard Foundation and University of Cambridge Climate Resilience and Sustainability Research Fund.

## DATA AVAILABILITY STATEMENT

This study analysed publicly available datasets; details of the sources and class compositions are provided in Table 1. Code, image embeddings, metadata, and model weights are available at github.com/Geofly/vision-normalizing-flows-for-banana-diseases.

## Notes

https://github.com/Geofly/vision-normalizing-flows-for-banana-diseases}

### Competing Interest Statement

The authors have declared no competing interest.

